# *MS1/MMD1* homologs in the moss *P. patens* are required for male and female gametogenesis and likely for sporogenesis

**DOI:** 10.1101/2022.03.27.485980

**Authors:** Katarina Landberg, Mauricio Lopez-Obando, Victoria Sanchez Vera, Eva Sundberg, Mattias Thelander

**Author notes:** Shared first authors (Katarina Landberg, Mauricio Lopez-Obando). Correspondence: Mattias Thelander.

## Abstract

- The Arabidopsis Plant HomeoDomain (PHD) proteins AtMS1 and AtMMD1 provide chromatin-mediated transcriptional regulation essential for tapetum-dependent pollen formation. Such pollen-based male gametogenesis is a derived trait of seed plants. Male gametogenesis in the common ancestors of land plants is instead likely to have been reminiscent of that in extant bryophytes where flagellated sperms are produced by an elaborate gametophyte generation. Still, also bryophytes possess MS1/MMD1-related PHD proteins.
- We addressed the function of two MS1/MMD1-homologs in the bryophyte model moss *Physcomitrium patens* by the generation and analysis of reporter and loss-of-function lines.
- The two genes are together essential for both male and female fertility by providing cell autonomous functions in the gamete-producing inner cells of antheridia and archegonia. They are furthermore expressed in the diploid sporophyte generation suggesting a function during sporogenesis, a process proposed related by descent to pollen formation in angiosperms.
- We propose that the moss MS1/MMD1-related regulatory network required for completion of male and female gametogenesis and possibly for sporogenesis, represent a heritage from ancestral land plants.

## Introduction

The Plant HomeoDomain (PHD) motif defines a family of proteins that can recognize and bind histones depending on covalent modification status of the histone tales (Mouriz et al., 2015). By recruitment and regulation of chromatin remodeling factors and transcriptional regulators, PHD proteins can thereby control chromatin compaction and gene expression in a histone modification-governed manner.

Phylogenetic analysis places angiosperm PHD proteins into five main subfamilies and a number of clades (Cao *et al*., 2018). Among these, clade IIa comprises members from both mono- and dicotyledonous species including the Arabidopsis genes *MALE STERILITY 1 (AtMS1*) and *MALE MEIOCYTE DEATH 1* (*AtMMD1*). *AtMS1* and *AtMMD1* encode similar protein products which are both essential for pollen production in anthers. Still, the two genes exert their functions in distinct anther cell types. Thus, *AtMMD1* controls gene expression and chromosome condensation needed for completion of meiosis in microsporocytes (Reddy *et al*., 2003; Yang *et al*., 2003) while *AtMS1* controls gene expression and function of tapetal cells surrounding and nursing the microsporocytes and microspores on their route towards functional pollen (Wilson *et al*., 2001; Ito and Shinozaki, 2002; Alves-Ferreira *et al*., 2007; Yang *et al*., 2007; Reimegård *et al*., 2017; Lu *et al*., 2020). Both genes exert their functions through modification of chromatin structure. Thus, AtMS1 activates genes organized in clusters by relaxation of chromatin condensation (Reimergård *et al*., 2017) and AtMMD1 can bind histone tails in a modification-dependent manner (Andreuzza *et al*., 2015; Wang *et al*., 2016). A detailed mode of action was recently proposed for AtMMD1 in meiotic cells where the protein is recruited to H3K4me3 marks allowing it to modulate the target specificity of nearby JUMONJI 16 (JMJ16) histone demethylases through a physical interaction dependent on its central MMD domain (Wang *et al*., 2020).

The developmental process controlled by *AtMS1* and *AtMMD1*, i.e. tapetum-assisted microspore and pollen formation facilitating downstream male gametogenesis, is a derived trait of angiosperms (Hackenberg & Twell, 2019). Gametogenesis in the common ancestors of all extant land plants is instead likely to have been reminiscent of that in bryophytes of today, where eggs and flagellated sperms are produced by female archegonia and male antheridia formed by a dominant haploid gametophyte generation (Renzaglia *et al*., 2000; Hackenberg & Twell, 2019). These gametophytic reproductive organs were eventually lost from the angiosperm lineage as part of a drastic reduction of the haploid gametophyte generation accompanied by increased complexity of the diploid sporophyte generation (Harrison, 2017). As part of this transition, pollen is hypothesized to have evolved from walled spores reminiscent of those in extant bryophytes thanks to two key evolutionary adaptations (Hackenberg and Twell, 2019). First, divisions in the male gametophyte generation were almost completely abolished to arrive at the situation in present-day angiosperms where only two sequential specialized divisions produce a pair of male gametes inside a vegetative cell from the primary meiotic product (the microspore). Second, breakage of the spore wall was deferred so that the cell divisions producing the two gametes could be completed within a still intact wall, today recognized as the angiosperm pollen wall. The proposed evolutionary origin of pollen gives that tapetum-derived pollen production in angiosperms is related by descent to tapetum-dependent spore formation in bryophytes (Lopez-Obando *et al*., 2022).

The existence of PHD clade IIa homologs also in gametophyte dominant bryophytes (Higo *et al*., 2016; Sanchez-Vera *et al*., 2022), separated from angiosperms for about 450 million years (Morris *et al*., 2018), suggest that clade IIa-related genes were present already in the common ancestors of all extant land plants. Recent transcriptome data from the bryophyte model moss *Physcomitrium patens* reports the expression of two clade IIa homologs in sporophytes at stages during which tapetal-like cells are active and spores and their precursors develop (Perroud *et al*., 2018; Lopez-Obando *et al*., 2022). Moreover, expression of the two genes was also detected in antheridia (male reproductive organs) and in the egg cell in archegonia (female reproductive organs), both produced by the haploid gametophytic generation (Meyberg *et al*., 2020; Sanchez-Vera *et al*., 2022 and references therein). Similarly, a putative clade IIa homologue of the model liverwort *Marchantia polymorpha* is also active in reproductive organs, at least in the antheridia (Higo *et al*., 2016). This points towards a function for bryophyte clade IIa genes during gametogenesis. Further dissection of this function may add to our understanding about the mechanisms, regulation and evolution of gametogenesis in land plants (Berger and Twell, 2011; Hisanaga *et al*., 2019).

Here we describe the functional characterization of the clade IIa PHD homologs *PpMS1A* and *PpMS1B* in the moss *P. patens. PpMS1A* and *PpMS1B* are together required for male and female fertility by providing cell autonomous functions essential for development of the gamete-producing inner cells of both antheridia and archegonia. The expression domains of the two genes furthermore suggest functions in sporogenous cells and in foot transfer cells of the diploid sporophyte generation. Based on these findings, we discuss a possible ancestral function for clade II PHD proteins and elaborate on how this may have evolved into the functions evident in present-day bryophytes and angiosperms, respectively.

## Materials and methods

### Plant material, growth conditions, tissue harvest, transformation and crosses

*Physcomitrium patens* (previously *Physcomitrella patens*) ecotype Reute (R) (Hiss *et al*., 2017) was used as WT and is the background to all transgenic lines in this study. Protonemal moss tissue was grown aseptically on solid BCD medium (Thelander *et al*., 2007) supplemented with 5 mM Ammonium Tartrate and 0.8% agar in petri dishes at 25°C under constant white light from fluorescent tubes (Philips F25T8/TL741, www.lighting.philips.com) at 35 μmol m^-2^s^-1^ in a Percival Scientific CU-41L4 growth chamber (www.percival-scientific.com). To induce reproductive organs and subsequent sporophyte development, young chloronemal tissue was shaped into round balls and placed on solid BCD medium in 15 mm deep petri dishes (90 mm in diameter). The ball-shaped tissue was allowed to grow out into gametophore-containing colonies for 5-6 weeks where after the plates were transferred to SD conditions (8 h of light, 30 μmol m^-2^s^-1^) at 15° C in a Sanyo MLR-350 light chamber to induce reproductive development. To enhance fertilization the plants were submerged in water overnight at 20±1 dpi. Crosses were carried out as described in Thelander *et al*. (2019) and the *P. patens* ecoype Gransden was used as a WT line with strongly reduced male fertility. For expression and phenotype analysis, gametophyte shoots harboring either reproductive organs or a developing sporophyte in the apex were harvested from the periphery of moss colonies at indicated time points. Under a Leica MZ16 stereo microscope (Leica Biosystems, Heidelberg, Germany), all leaves were removed to expose the antheridia and archegonia. For sporophyte analysis also residual reproductive organs were detached, as were the sporophyte calyptra from stage 8. To enhance penetration, sporangia harvested after 12 dpw were punctuated using a fine needle. Protoplast transformation was carried out as previously described (Schaefer *et al*., 1991). Stable transformants were selected in the presence of 50 μgml^-1^ hygromycin (Duchefa H0192; Haarlem, the Netherlands) or G418 (11811023; Thermo Fisher Scientific, Waltham, MA, USA).

### Generation of reporter lines

Primer sequences are shown in Table S1. The *PpMS1A* translational reporter construct pMLO14 (Fig. S1a), used to integrate a GFP-GUS gene in frame near the end of the coding sequence was generated by the fusion of four PCR fragments using In-Fusion technology (www.takarabio.com): A 4550 bp vector fragment amplified with primers SS748/SS749 from plasmid pDEST14 (www.thermofisher.com), a fragment covering 669 bp from exon 3 to near the end of the *PpMS1A* CDS amplified with primers SS750/SS751 from WT gDNA, a fragment covering a 2556 bp GFP-GUS gene amplified with primers SS752/SS753 from plasmid pMT211 (Thelander *et al*., 2019), and a fragment covering 651 bp of the extreme end of the CDS and the 3’UTR of *PpMS1A* amplified with primers SS754/SS755 from WT gDNA. To generate the *PpMS1Apro::PpMS1A-GFPGUS* reporter lines, 8 μg of pMLO14 and 4 μg of pMLO13 were co-transformed into WT protoplast together with 8 μg pACT1:hCAS9 and 4 μg of pBNRF (Lopez-Obando *et al*., 2016). Stable transformants were selected on G418, where after the in frame-fusion between the *PpMS1A* CDS and the GFP-GUS gene resulting from correct integration was confirmed by PCR amplification using the primers SS756/SS627 followed by sequencing of the resulting PCR product with the primers SS757 and SS627. Three independent lines showing correct integration were selected for downstream analysis (Table S2). The three lines indicated qualitatively similar signal patterns, but while signals in *PpMS1Apro::PpMS1A-GFPGUS-1* were strong and coherent in both reproductive organs and sporophytes, signals in *PpMS1Apro::PpMS1A-GFPGUS-2* and *-3* were generally weaker, and, as a consequence of this, challenging to detect in reproductive organs.

To produce the *PpMS1B* transcriptional reporter construct pVS1 (Fig. S1b), the *PpMS1B* promoter was amplified from gDNA with primers SS738/SS739, trimmed to 2918 bp with *BamHI/NcoI*, and cloned between the same sites of the vector pMT211. The vector pMT211 carries a hygromycin selection cassette and allows promoters to be cloned ahead of a *GFP-GUS* reporter gene for subsequent integration into the *Pp108* locus (Thelander *et al*., 2019). The resulting construct was verified by sequencing and linearized with *Sfi*I before transformation into WT moss. Correct integration was confirmed by PCR-verification of 5’ and 3’ junctions with the primers SS5/SS742 and SS399/SS307, respectively Fig. S1c,d). Three independent lines showing correct integration (*PpMS1B::GFPGUS-1,2,3*) were selected for downstream analysis and were found to display essentially identical reporter signals in all tissues investigated.

### Generation of loss-of-function mutants

Primer sequences are shown in Table S1. *PpMS1A* loss-of-function mutants were generated by CRISPR technology, and gRNA-expressing constructs were designed using CRISPOR (Haeussler *et al*., 2016). For each gRNA used putative off-targets had at least four mismatches making off-target editing events highly unlikely (Table S3; Modrzejewski *et al*., 2020). To produce the plasmids pMLO11 and pMLO12 (Table S3; Fig. S2a), AttB1-PpU6-SgRNAs-AttB2 fragments produced by gene synthesis (Integrated DNA Technologies, Coralville, USA) were cloned into the vector pDONR221 by Gateway recombination (Invitrogen, Carlsbad, USA). To produce plasmid pMLO13 (Table S3; Fig. S2a), the annealing product of the complementary primers SS743/SS744 was cloned into the vector pENTR_PpU6_L1L2 opened with *BsaI* (Mallett *et al*., 2019). Inserts were confirmed by sequencing. CRISPR mutants were then obtained as previously described (Lopez-Obando *et al*. 2016). In short, WT or *ms1b-1* (see below) protoplasts were co-transformed with 8 μg of pACT1:hCAS9, 4 μg of pBNRF, and 4 μg of each of the plasmids pMLO11 and pMLO12 or pMLO13. Transformants were selected in presence of G418 and mutations were evaluated by PCR amplification and sequencing of gDNA with the gene specific primers SS745/SS746 and SS745/SS747 (Table S2; Fig. S2a). For phenotypic analysis, three single (*ms1a*-*1*,*2*,*3*) and two double (*ms1ams1b-1*,*2*) mutant lines with mutations in *PpMS1A* likely to block protein function were selected for phenotypic analysis (Table S2). Independent lines of the same genotype were found to display essentially identical phenotypes in all tissues examined.

*PpMS1B* loss-of-function mutants were generated by homologous recombination, and to produce the *PpMS1B* knockout construct pVS2 (Fig. S2b), Gateway 3-fragment recombination technology was used (www.thermofisher.com). Thus, LR recombination was used to fuse a 909 bp *PpMS1B* 5’ fragment (amplified with the primers SS734/SS735 and cloned into the entry vector pDONR P1-P4), a G418 resistance fragment from the entry clone pDONR4r-3r-G418 (Landberg *et al*., 2020), and a 1180 bp *PpMS1B* 3’ fragment (amplified with the primers SS736/SS737 and cloned into the entry vector pDONR P3-P2), into the destination vector pDEST14. The resulting construct was verified by sequencing and was linearized with *HpaI/AvrI* before transformation into WT moss and the selection of stable transformants in presence of hygomycin. Correct integration was confirmed by PCR-verification of 5’ and 3’ junctions with the primers SS58/SS759 and SS762/SS763, respectively (Fig. S2c). Two independent lines showing correct integration (*ms1b-1*,*2*) were selected for downstream analysis and were found to display essentially identical phenotypes in all tissues investigated.

### RT-qPCR

For the analysis of *PpMS1A* and *PpMS1B* expression in various WT tissues, samples from gametophore apices harvested at different time points after induction, antheridia, archegonia, and sporophyte samples corresponding to different developmental stages have been previously described (Landberg *et al*., 2020; Lopez-Obando *et al*., 2022). Tissue harvest, RNA extraction, cDNA synthesis and amplification, setup and cycling of qPCR reactions, normalization using three reference genes, and calculations have been previously described (Landberg *et al*., 2020). The gene-specific primers used were SS586/SS587 for *PpMS1A* and SS584/SS585 for *PpMS1B* (Table S1). To avoid amplification of genomic DNA contaminations, the annealing site for one primer in each pair is interrupted by an intron. Melt curve, gel and standard curve analyses confirmed that both primer pairs amplified a single product of the expected size with efficiencies close to 100 % (data not shown). Data is presented as relative expression calculated with the 2–ΔΔCT method. In Fig. 1c-d the sample with the highest transcript abundance for each gene was set to 1. In Fig. S3a-b, the same data is presented, but here the sample with the highest overall transcript abundance (regardless of whether it was *PpMS1A* or *PpMS1B*) was set to 1. Each data point is based on biological triplicates and error bars represent standard deviations.

**Figure 1.**
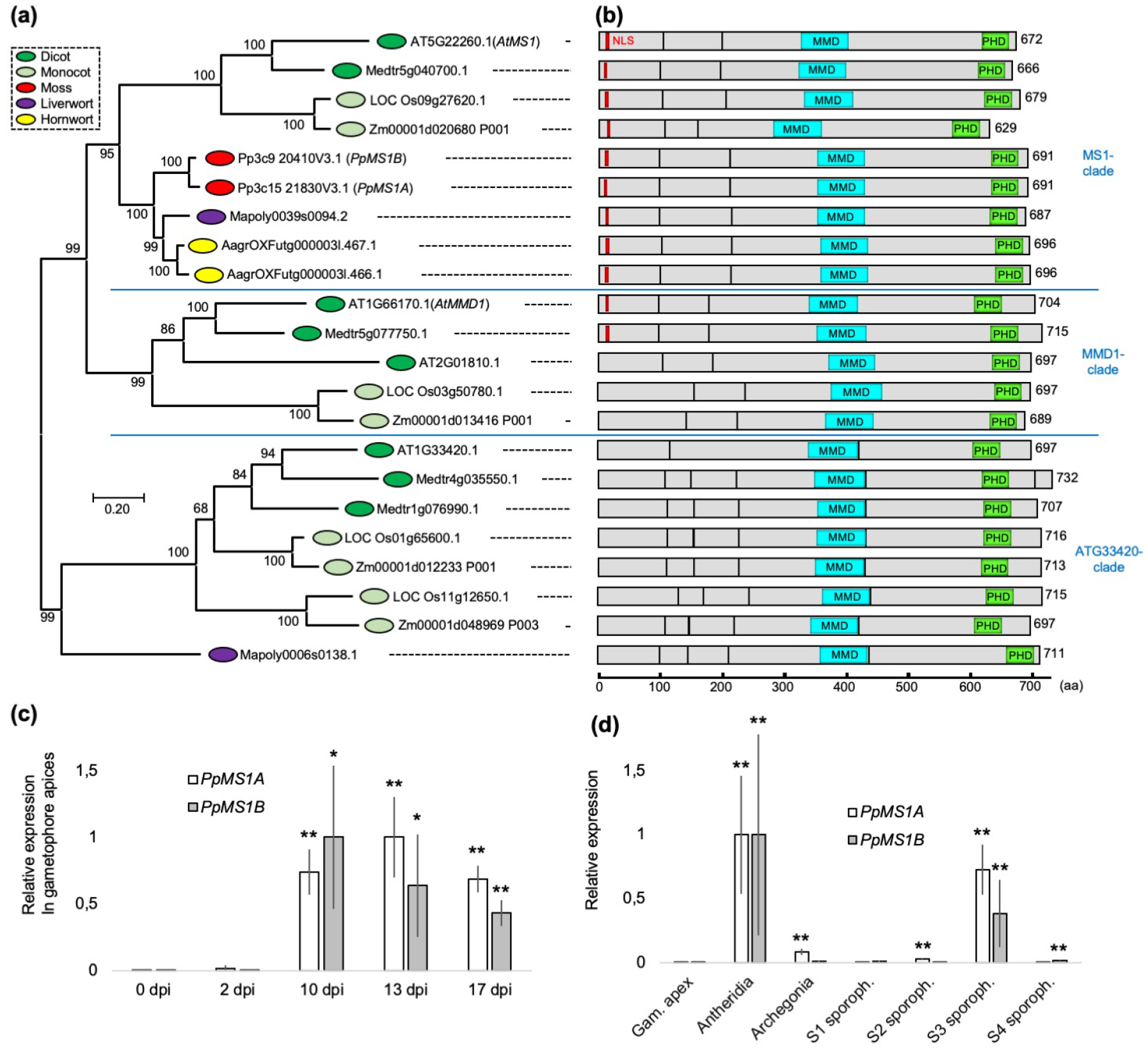
Phylogeny of PHD clade IIa bryophyte homologs and gene expression in the moss *P. patens* in sporophytes and reproductive organs. (a) Maximum likelihood tree demonstrating the phylogenetic relation of PHD clade IIa-related transcription factors from selected angiosperm and bryophyte species. AT, *Arabidopsis thaliana;* Medtr, *Medicago truncatula;* LOC Os, *Oriza sativa;* Zm, *Zea mays;* Pp, *Physcomitrium patens;* AagrOXF, *Anthoceros agrestis;* Mapoly, *Marchantia polymorpha*. (b) Schematic view of proteins from (a) demonstrating exon organization and the positions of conserved domains that have been assigned possible functions in flowering plants. Gray box, exon; red box, putative nuclear localization signal consisting of four or more basic amino acids; blue box, MMD protein interaction domain; green box, PHD domain. For a full length amino acid alignment of proteins belonging to the MS1- and the MMD1-clades with the same domains marked, see Fig. S5. (c) Relative transcript abundance of *PpMS1A* and *PpMS1B* in WT gametophore apices at different days post induction (dpi) of reproductive development. Typical occurrence of reproductive organs at selected time points: 0 and 2 dpi, no reproductive organs; 10 dpi, young antheridia; 13 dpi, mid-stage antheridia and young archegonia; 17 dpi, mature antheridia and mid-stage archegonia. (d) Relative transcript abundance of *PpMS1A* and *PpMS1B* in isolated WT gametophore shoot apices (without reproductive organs), antheridia bundles, archegonia bundles, and sporophytes of different developmental stages. Sporophyte samples contain organs roughly correlating to the following stages described in Lopez-Obando *et al*. (2022): S1, st.1-3; S2, st.4-8; S3, st.9-11; S4, st.12-14. In both (c) and (d), the sample with the highest transcript abundance for each gene is set to 1, each data point represents an average of three independent biological replicates, error bars indicate standard deviation and asterisks indicate a statistically significant difference from gametophore apex sample prior to reproductive organ formation (Student’s t-test: *, P < 0.05; **, P < 0.02). See also Figure S3a-b for presentation of the same data in a way making comparisons of transcript abundance levels between *PpMS1A* and *PpMS1B* possible.

### Sequence retrieval, proteins alignments and phylogenetic analysis

Gene and protein sequences of PHD clade IIa homologs displayed in the phylogenetic tree of Fig. 1A were retrieved from Phytozome V12.1 and www.hornworts.uzh.ch following BLAST-based gene identification. For the phylogenetic reconstruction, amino acid sequences were aligned using the M-Coffee algorithm in T-Coffee (Notredame *et al*., 2000; Wallace *et al*., 2006) where after the alignment was filtered using Transitive Consistency Scores (Chang *et al*., 2014). The resulting filtered alignment (Fig. S4) was used for phylogenetic reconstruction in Megax (v.10.1.5; Kumar *et al*., 2018) with the maximum likelihood method (JTT amino acid substitution model, gamma distribution among sites) and 500 replications of bootstrapping. The non-filtered alignment of full length MS1- and MMD1-clade proteins in Fig. S5 was produced with the MUSCLE algorithm in Megax (v.10.1.5; Kumar *et al*., 2018). Alignments in Fig. S4 and S5 were displayed with AliView (Larsson, 2014).

### GUS staining

Gametophore apices with reproductive organs or sporophytes were harvested as described above and incubated in GUS solution (50mM NaPO4, pH 7.2, 2 mM Fe2+CN, 2 mM Fe3+CN, 2 mM X-Gluc and 0.2% (v/v) Triton X-100) at room temperature for 48 h. For analysis of intact organs the tissue was transferred to 70% EtOH and prior to analysis the organs were mounted on objective glasses in 30 % glycerol. For sectioning, the tissue was transferred from GUS solution to FGPX fixative (Lopez-Obando *et al*., 2022) and then treated as describe below.

### Sporophyte sectioning

Thin sectioning of sporophytes was carried out as described in Lopez-Obando *et al*. (2022).

### Microscopy

Intact GUS-stained reproductive organs and sporophytes mounted in 30 % glycerol as well as Toulidine blue- or GUS-stained sections of mutant and reporter lines were analysed using an Axioscope A1 microscope equiped with an AxioCam ICc 5 camera and the Zen Blue software (Zeiss) at 10x, 20x and 63x magnification. Images of mutant antheridia and archegonia mounted in 30 % glycerol were captured using a DMI4000B microscope with differential interference contrasts optics at 63x magnification, a DFC360FX camera, and the LAS AF software (Leica microsystems). For GFP expression analysis, reproductive organs and developing sporophytes of indicated reporter lines were harvested and mounted in water immediately prior to analysis. Signals were documented using a LSM 780 confocal laser scanning microscope (Carl Zeiss) with a GaAsP detector and 20x (NA 0.8) and 63x (NA1.2, water immersion) objectives. Excitation/detection parameters were 488 nm/491-598 nm for GFP and 633/647-721 nm for chlorophyll auto-fluorescence. Images were acquired using ZEN black software and are snapshots of a single focal plane with selected channels overlaid. Adobe Photoshop CC was used to adjust intensity and contrast, mark borders and cells, cut away surrounding areas, and merge images to visualize entire large organs at high magnification. Bar charts, tables, calculation of means, standard deviation and Student t-tests were performed using Microsoft Excel.

## Results

### The emergence of PHD clade IIa transcription factors predated the divergence of bryophytes and angiosperms

Genome-wide phylogenetic analyses of angiosperm PHD genes indicate that *AtMS1* and *AtMMD1* (and its close homolog AT2G01810.1) emerged through a duplication event predating the evolutionary split into mono- and dicots (Cao *et al*., 2018). Putative clade IIa PHD genes also exist in bryophytes even if their phylogenetic position is unknown (Higo *et al*., 2016; Sanchez-Vera *et al*., 2022). To change this, we screened genomes from representatives of land plant lineages for PHD clade IIa genes and subjected deduced amino acid sequences to phylogenetic analyses. This confirmed the existence of clear PHD clade IIa genes in all three main lineages of bryophytes, *i.e*. mosses, liverworts, and hornworts. All three lineages have genes clustering within the angiosperm *MS1*-subclade but lack genes clustering within the *MMD1*-subclade (Fig. 1a, S4). The liverwort *M. polymorpha* also has a gene clustering within the subclade of AT1G33420.

*PHD* clade IIa genes in bryophytes and angiosperms furthermore share the same exon/intron-organization and encode proteins with similar domain structure (Fig. 1b). Striking sequence similarity is evident throughout large parts of the proteins, and is particularly pronounced in regions demonstrated to be functionally important in angiosperm MS1 and/or MMD1 proteins. Thus, conserved regions include the C-terminal PHD domain, a suggested N-terminal nuclear localization signal, and the internal MMD domain which in AtMMD1 regulates the substrate specificity of the JMJ16 histone demethylase by a physical interaction (Fig. 1b, S5; Wilson *et al*., 2001; Ito and Shinozaki *et al*., 2002; Reddy *et al*., 2003; Yang *et al*., 2003; Wang *et al*., 2020).

### *PpMS1A* and *PpMS1B* are expressed in developing sporophytes, as well as in male and female reproductive organs

To reveal the function of PHD clade IIa genes in bryophytes, and with hope of gaining insight about ancestral functions of this gene family in land plants, we have functionally characterized the two *Physcomitrium patens* clade IIa homologs. Based on their clustering with genes belonging to the *MS1*-subclade (Fig. 1a), we call the genes *PpMS1A* (Pp3c15_21830V3.1) and *PpMS1B* (Pp3c9_20410V3.1). As published transcriptome data indicates that *PpMS1A* and *PpMS1B* are exclusively expressed in the early sporophyte generation, and in reproductive organs produced by the haploid gametophyte generation (Sanchez-Vera *et al*., 2022 and references therein), we first verified this using qPCR. We checked expression in gametophore (gametophytic shoot) apices at different times post induction (dpi) of reproductive development (for typical ontogeny, see Landberg *et al*., 2013). This revealed a complete lack of expression in apices yet to develop reproductive organs (0, 2 dpi), but clear expression of both genes in apices which had developed reproductive organs (10, 13, 17 dpi) (Fig. 1c, S3a). Next, we checked expression in isolated antheridia and archegonia, as well as in isolated sporophytes of different developmental stages. This revealed significantly increased expression in all three tissue types of at least one of the genes when compared to vegetative shoot apices (Fig. 1d, S3b). Antheridia showed highly significant expression of both genes while expression in archegonia was lower and proved significant only for *PpMS1A*. The relatively low expression in archegonia may be explained by expression in only a fraction of the cells in the organ, a prospect fitting well with the previous observation that expression of the two genes is scored in eggs but not in cavity wall cells (Sanchez-Vera *et al*., 2022). In sporophytes, transcript abundance of both *PpMS1A* and *PpMS1B* peaked in the S3 sample, corresponding to developmental stages 9-11 in Lopez-Obando *et al*. (2022), indicating active expression sometime between completion of embryogenesis and the appearance of mature spores. Finally, even if comparisons of expression between genes based on qPCR data should be handled with care, our data indicates that *PpMS1A* is expressed at generally higher levels than *PpMS1B* (Fig. S3a, S3b).

### Cell autonomous *PpMS1* activity is essential for developmental progression of spermatogenous cells

In the haploid gametophyte generation, all three major lineages of bryophytes produce flagellated sperms in antheridia (Renzaglia *et al*., 2000). In *P. patens*, the vegetative shoot apex is reprogrammed to produce antheridia in response to low temperatures and short day-length through a stereotypic developmental program (Hohe *et al*., 2002; Landberg *et al*., 2013; Hiss *et al*., 2017; Kofuji *et al*., 2018). To get a more detailed picture of where and when during antheridia development the *PpMS1* genes are expressed, we produced translational reporter lines for *PpMS1A (PpMS1Apro::PpMS1A-GFPGUS-1*,*2*,*3*) and transcriptional reporter lines for *PpMS1B* (*PpMS1Bpro::GFPGUS-1*,*2*,*3*) (Materials and Methods; Fig. S1).

The translational *PpMS1A* reporter showed signals in antheridial inner cells from stage 3 to 7, with a peak around stage 5, while no signals were detected in jacket and tip cells (Fig. 2a; stages defined in Landberg *et al*., 2013). PpMS1A expression is thus restricted to the spermatogenous cells, comes on as soon as they appear, stays active throughout their division phase, and fades out at around their entrance into spermatogenesis. In contrast, the transcriptional *PpMS1B* reporter failed to detect expression in any stage or in any part of antheridia. This fits well with our qPCR data indicating a significant but manifold lower expression of *PpMS1B* than of *PpMS1A* in antheridia (Fig. S3a,b).

**Figure 2.**
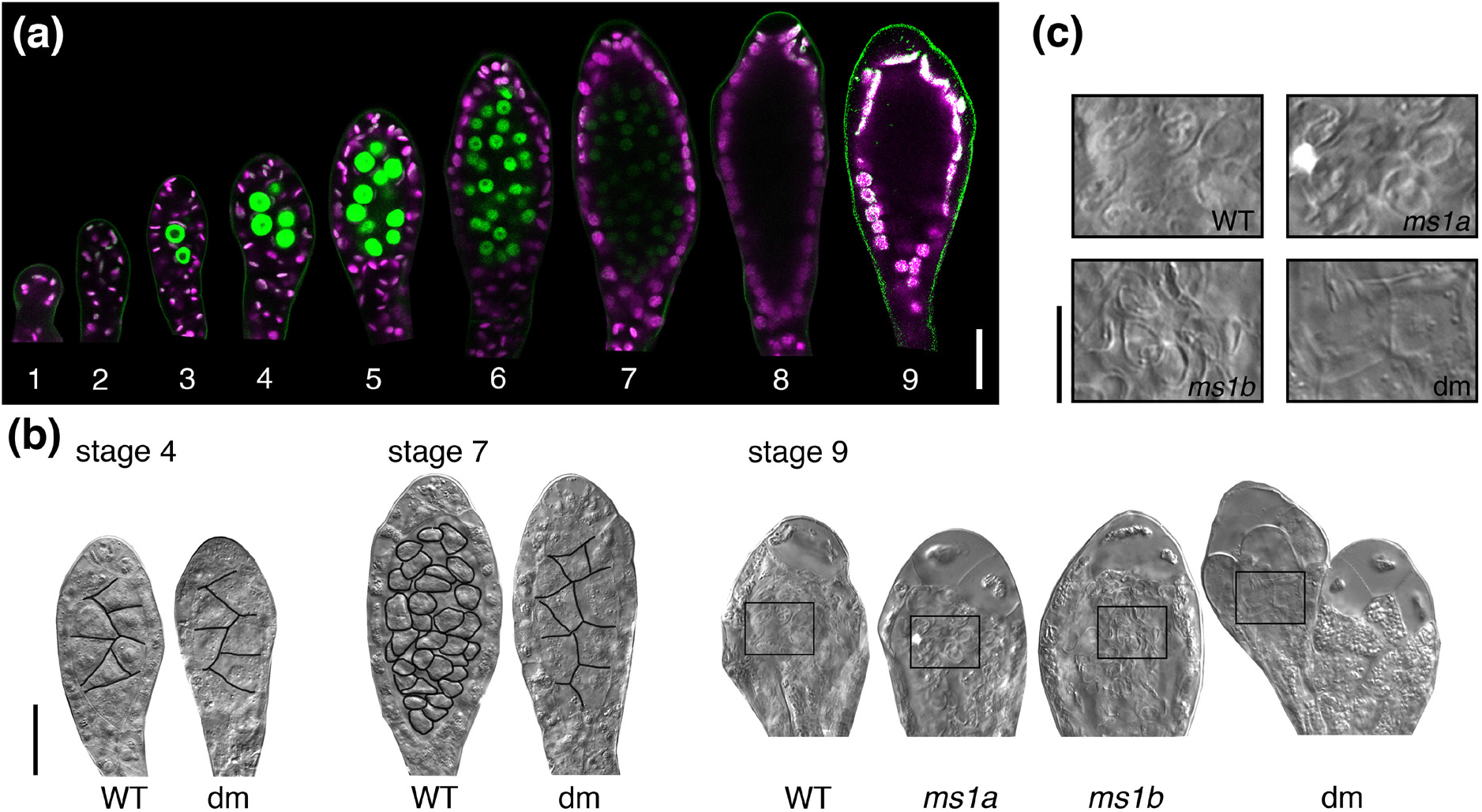
*PpMS1* functions in antheridia. (a) *PpMS1A::PpMS1A-GFPGUS-1* reporter signals in representative antheridia. A merge of confocal channels detecting green fluorescent protein (green) and chloroplast autofluorescence (magenta) are shown, and the numbers 1-9 indicate developmental stages according to Landberg *et al*. (2013). Note expression in spermatogenous inner cells from stage 3 to 7. Bar, 20 μm. (b) Differential interference contrast images of representative stage 4, 7, and 9 antheridia from WT and the *ms1ams1b-1* double mutant (dm). Note problems with proliferation and differentiation of spermatogenous inner cells in dm. Stage 9 *ms1a-1* and *ms1b-1* single mutant antheridia are also shown to demonstrate normal sperm production in these genotypes. Borders between spermatogenous cells have been traced in black for clarity. Bar, 20 μm. (c) High magnification of boxed areas in stage 9 organs in (b). Bar, 10 μm.

To address the functional relevance of *PpMS1* expression in antheridia, we went on to produce single and double loss-of-function mutants for the two genes. *PpMS1A* single mutants (*ms1a-1*,*2*,*3*) were produced by CRISPR editing, *PpMS1B* single mutants (*ms1b-1*,*2*) were produced by homologous recombination, and double mutants (*ms1ams1b-1*,*2*) were produced by CRISPR editing of *PpMS1A* in the *ms1b-1* background (Materials and Methods; Fig. S2). Although neither single loss-of-function mutant showed obvious deviations in antheridia development, the male organs of *ms1ams1b* double mutants displayed an arrest of inner cell development resulting in a complete inability to produce functional sperms (Fig 2b,c). The periclinal divisions in the early antheridium giving rise to the first 4-6 inner cells appeared unaffected, and the newly formed inner cells typically also divided once, but after this, inner cell divisions ceased in the double mutant. WT inner cells continue to divide on the expense of cell size during stages 4-6 where after they enter spermatogenesis at stage 7, but double mutant inner cells instead remained large and kept an appearance similar to the outer cells from which they first originated. The sterile jacket and tip cells of double mutant antheridia matured as in WT even if the organ tip failed to open at maturity. This suggests that jacket and tip cell maturation is independent on successful differentiation of inner cells into sperms, but that the bursting of organ tips may somehow require the formation of sperms.

### Cell autonomous *PpMS1* activity is needed for canal clearance and egg cell maturation in archegonia

*P. patens* female reproductive organs (archegonia) are initiated from a lateral position close to the gametophore apex a few days after the outgrowth of the first antheridia, and their development has been described previously (Kofuji *et al*., 2009; Landberg *et al*., 2013; 2020). Signals in archegonia from the translational *PpMS1A* reporter were restricted to the central cell file, consisting of a basal-most pre-egg/egg, an upper basal cell, and four apical canal cells (Fig. 3a). The signals indicated relatively strong PpMS1A protein expression in the pre-egg/egg and the upper basal cell from stage 5 to 7, where after expression in these two cells declined successively during stage 8 and 9, when the egg matures and the canal cells degrade (stages defined in Landberg *et al*., 2013). The reporter also indicated weak PpMS1A expression in canal cells from stage 6 to 8. As in antheridia, we were unable to detect signals from the transcriptional *PpMS1B* reporter in archegonia, probably reflecting generally low expression levels as indicated by our initial qPCR experiments (Fig. 1d, S3d).

**Figure 3.**
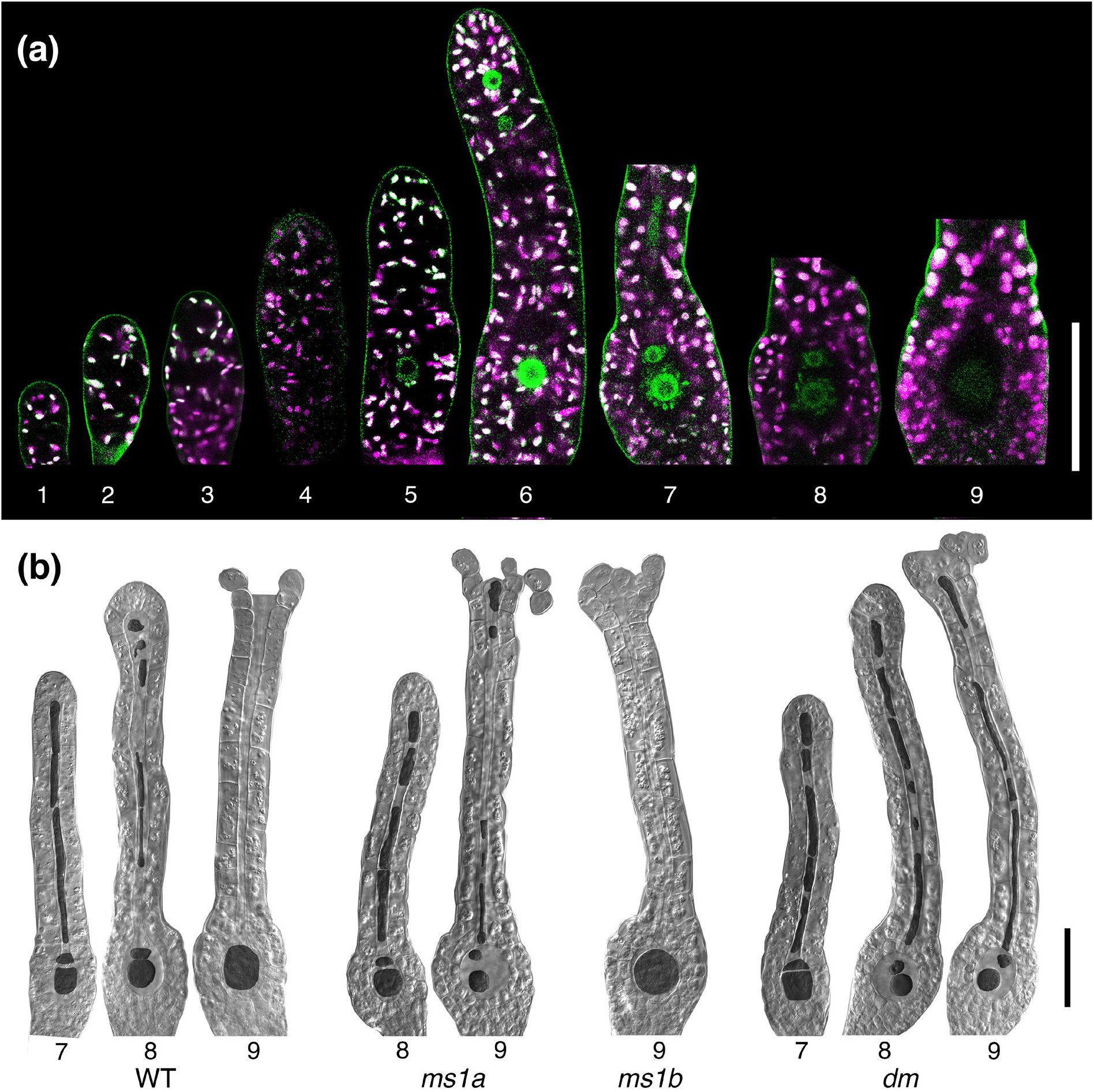
*PpMS1* functions in archegonia. (a) *PpMS1A::PpMS1A-GFPGUS-1* reporter signals in representative antheridia. A merge of confocal channels detecting green fluorescent protein (green) and chloroplast autofluorescence (magenta) are shown, and the numbers 1-9 indicate developmental stages according to Landberg *et al*. (2013). Note expression in central file of inner cells. Bar, 50 μm. (b) Differential interference contrast images of representative stage 7, 8, and 9 antheridia from WT and the *ms1ams1b-1* double mutant (dm). Note problems with degradation of upper basal cell and canals cells during stage 8 and 9, and the reduced size of the egg at stage 9 in dm. Stage 8 and 9 *ms1a-1* organs, displaying a mild version of the dm phenotype, and stage 9 *ms1b-1* single mutant archegonia, displaying no clear phenotype, are also shown. Inner cells have been false colored in dark grey for clarity. Bar, 50 μm.

The phenotypic defects of the *ms1ams1b* archegonia correlates spatially and temporally with the expression domains of the *PpMS1A* reporter (Fig. 3b). Archegonia from the *ms1ams1b* double mutant developed normally up until stage 7, by concluding cell divisions in the central cell file including an asymmetric division of the basal-most cell to produce the egg and the upper basal cell. However, they consistently failed to complete degradation of the upper basal cell and canal cells during stage 8, which is required for the formation of an open canal down to the egg through which sperms can enter when the organ tip opens at stage 9 (Fig. 3b). This deficiency prevented canal clearance, and is likely to block sperm access to the egg even if the organ tip opened as in WT during stage 9. We also observed that egg cells in stage 9 *ms1ams1b* archegonia were generally smaller and more compact than in WT (Fig. 3b).

While the *ms1b* single mutant produced WT-like archegonia, the female reproductive organs of the *ms1a* single mutant showed similar but milder deficiencies compared to *ms1ams1b* organs, with only partially blocked degradation of upper basal and canal cells, and lower penetrance of abnormal egg cell compaction (Fig. 3b). The fact that phenotypes in the *ms1ams1b* double mutant is more severe than in the *ms1a* single mutant suggests that *PpMS1B* contributes to *PpMS1* functions in archegonia, at least in the absence of *PpMS1A*.

### *PpMS1* functions are essential for both male and female fertility

*P. patens* is a monoecious species fully capable of self-fertilization (Perroud *et al*., 2019). To assess the effect of reduced or lost *PpMS1* function on fertility, we next carried out a series of crossing experiment. The results confirmed that the *ms1ams1b* double mutant is completely unable to self (Table 1). This was expected since the double mutant cannot produce sperms (see above), but crosses also showed that sporophyte production from mutant archegonia could not be restored even when the highly fertile Reute WT was used as the sperm donor, indicating that the double mutant is both male and female sterile (Table 1).

**Table 1.**
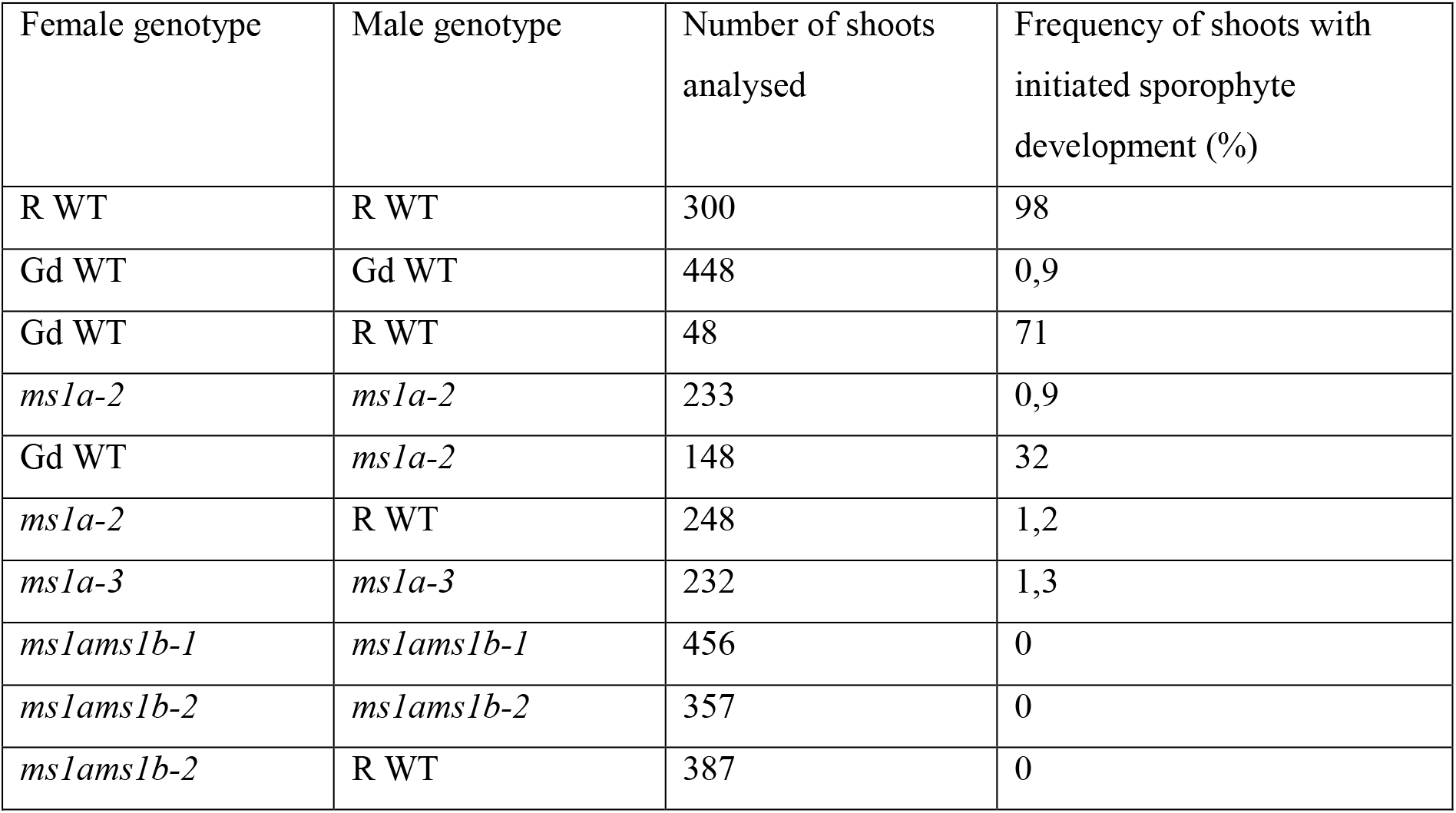
Fertility of *ms1* mutants. Number of shoots that formed sporophytes after selfing or after crosses between indicated *ms1* mutant line and wild-type (WT) strains Reute (R) or Gransden (Gd).

We also explored the fertility of the *ms1a* single mutant by subjecting it to selfing and crosses to both the fertile Reute WT and the largely male sterile Gransden WT (Landberg *et al*., 2020; Meyberg *et al*., 2020). The crosses showed that the *ms1a* single mutant can self, albeit at much lower frequencies than its parental Reute WT, indicating that the single mutant suffers from reduced fertility but is able to produce fertilization competent male and female gametes to some extent (Table 1). The much-reduced fertilization frequency could not be restored even when the highly fertile Reute WT was used as the sperm donor, indicating that the deficiency is mainly due to a female fertility problem (Table 1). This conclusion is partially supported by the ability of *ms1a* single mutant sperms to significantly increase the frequency of sporophyte formation from archegonia of the largely male sterile Gransden WT, demonstrating that *ms1a* single mutant sperms are largely functional (Table 1).

### *PpMS1* expression domains suggests functions in the foot and in sporogenous cells of the sporophyte

*PpMS1A* and *PpMS1B* are also active in mid-stage sporophytes supporting a *PpMS1-function* also in the diploid generation (Fig 1d, S3b). Like in all bryophytes, the moss zygote develops into a non-branched sporophyte consisting of a foot and an apical capsule (sporangium) in which sporogenous cells undergo meiosis to form spores. For an overview of *P. patens* sporophyte/sporangium ontogeny, including a definition of the developmental stages referred to below, see Lopez-Obando *et al*. (2022) and references therein.

Unfortunately, the PpMS1 loss-of-function mutants failed to provide clues to the functional relevance of *PpMS1* expression in sporophytes. As already described, the *ms1ams1b* double mutant was completely unable to produce sporophytes due to reproductive organ and gamete deficiencies (Fig. 2,3; Table 1), while the two single mutants produced WT-like sporophytes (albeit at reduced rates in the *ms1a* mutant) (Fig. 4a). This suggests that the two *PpMS1* genes have redundant functions during sporophyte development, a prospect supported by the fact that expression of the two genes in sporophytes shows temporal overlap and is more uniform in strength than in reproductive organs (Fig 1d, S3b). In addition, the reporter lines also revealed a spatial overlap of *PpMS1A* and *PpMS1B* expression in two discrete domains during sporophyte development. While no signals were detected in young embryos, both reporters were actively expressed in the foot of the slender embryo with an onset at around stage 5 (Fig. 4b, S6). The signals were largely concentrated to the epidermal transfer cells suggested to be important for nutrient uptake from the gametophore (Regmi & Gaxiola, 2017). Signals from the translational *PpMS1A* reporter eventually spread from the foot to parts of the seta connecting the foot to the sporangium, and largely persisted until sporophyte maturity. In contrast, signals from the transcriptional *PpMS1B* reporter were more restricted to the sporophyte foot and peaked in strength at around stage 8 where after they faded out.

**Figure 4.**
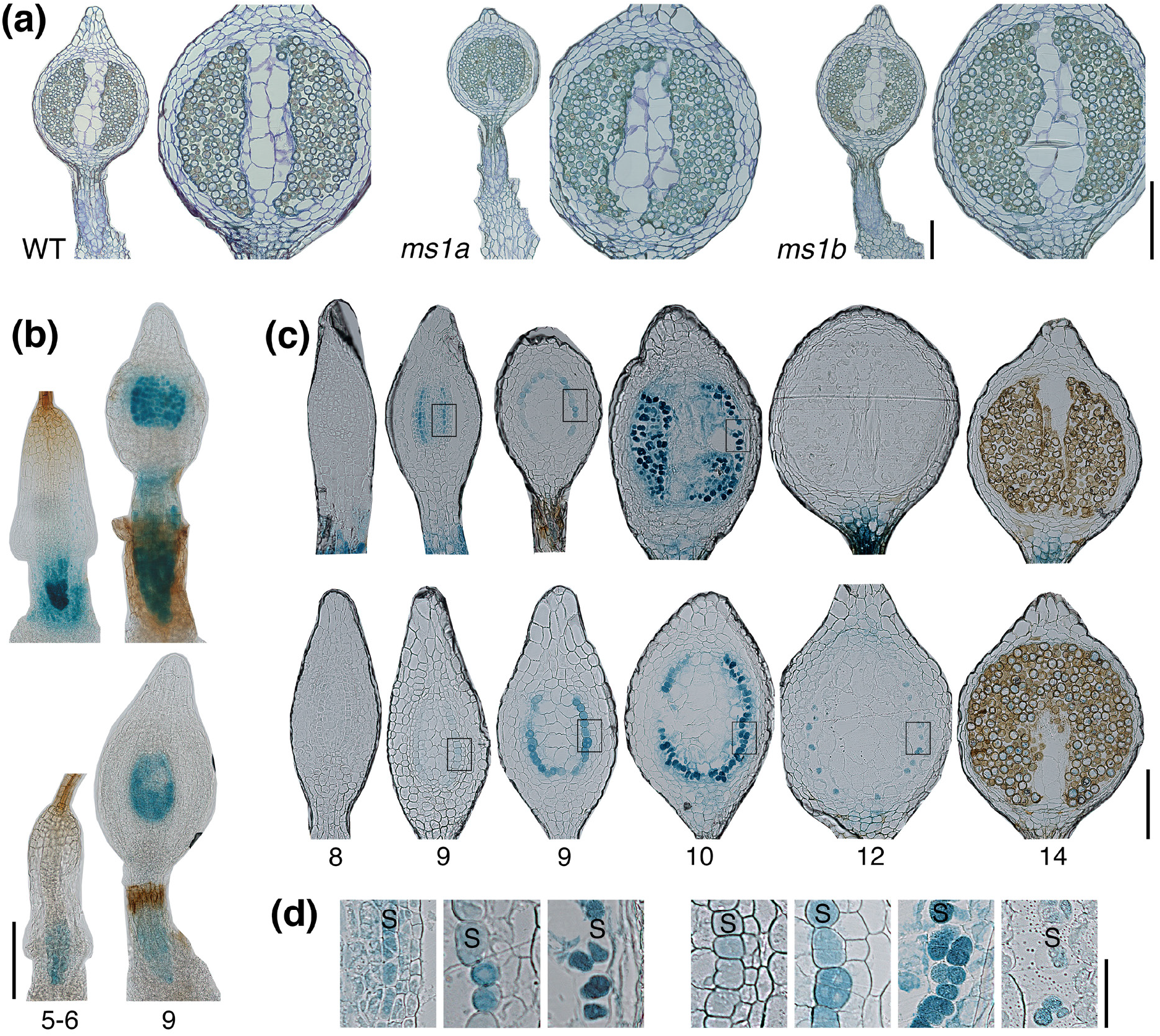
*PpMS1* activity in sporophytes. (a) Medial longitudinal sections through WT, *ms1a-2*, *ms1b-1* stage 14 sporophytes to demonstrate the lack of phenotypes in *ms1* single mutants. For each genotype, an entire sporophyte is shown to the left and a magnification of a sporangium to the right. (b-d) *PpMS1A::PpMS1A-GFPGUS-1* and *PpMS1B::GFPGUS-1* GUS reporter signals in sporophytes. (b) Whole-mounted stage 5-6 and 9 GUS-stained sporophytes of the *PpMS1A::PpMS1A-GFPGUS-1* (upper) and the *PpMS1B::GFPGUS-1* (lower) reporter lines. Note early signals in sporophyte foot and later signals central part of sporangium. (c) Sections of stage 8, 9, 10, 12 and 14 GUS-stained sporangia of the *PpMS1A::PpMS1A-GFPGUS-1* (upper) and the *PpMS1B::GFPGUS-1* (lower) reporter lines. Note that signals largely restricted to the sporogeneous cell layer comes on at stage 9. (d) High magnification of boxed areas in C. *PpMS1A::PpMS1A-GFPGUS-1* signals (left) are largely restricted to the sporogenous cell layer (S) even if a putative signal is evident also in the surrounding cells at early stage 9. *PpMS1B::GFPGUS-1* signals are restricted to the sporogenous cell layer (S) from the onset at stage 9. Numbers in (b) and (c) indicate developmental stages according to Lopez-Obando *et al*. (2022). See also Fig. S6 for *PpMS1B::GFPGUS-1* GFP reporter signals in sporophytes. Size bars in (a-c), 200 μm. Size bar in (d), 50 μm.

In addition to expression in the sporophyte foot, the two reporters also indicated expression of *PpMS1A* and *PpMS1B* in the sporogenous cells. Thus, signals from both reporters were evident in the sole sporogenous cell layer from early stage 9, *i.e*. immediately following completion of the cell division phase giving rise to the different cell layers of the sporangium (Fig. 4b-d, S6). The signals in the sporogenous cells persisted during their final division at stage 10 and their maturation into liberated globular sporocytes at stage 11, where after the signals started to fade out before meiosis and spore formation at stage 12 (Fig. 4c,d, S6). Signals from the translational *PpMS1A* reporter peaked at stage 9-10 and were completely lost around meiosis at stage 12 (Fig. 4c,d). Signals from the transcriptional *PpMS1B* reporter peaked at stage 10-11, dropped in intensity at around meiosis at stage 12, but were often weakly evident as late as in mature spores (Fig. 4c,d, S6). Signals from the two reporters were generally evident only in the sporogenous cells, but we came across a few examples of stage 9 sporophytes where the *PpMS1A* reporter showed weak putative signals also in the tapetum and columella layers, opening for the possibility that *PpMS1* activity could play a role also in these cells during a brief developmental window (Fig. 4c,d).

## Discussion

This study reveals that PHD clade IIa genes in moss control developmental processes in antheridia, archegonia and likely in sporophytes. *PpMS1* activity is absolutely essential both for male and female fertility, and fundamental features of how it affects the development of antheridia and archegonia are shared. *PpMS1* activity thus appears dispensable for the initiation of the two organ types, the development of their sterile structural parts, and the inwards formative divisions giving rise to the gamete-producing inner cell population. Instead, cell autonomous *PpMS1* activity is needed for the specification and further development of these inner cells. In antheridia, *PpMS1* controls the proliferation and differentiation of inner cells into sperms. In archegonia, *PpMS1* is needed for proper maturation of the egg, and for degradation of the remaining inner cells to leave a free canal passage for sperms to access the egg.

The reporter genes indicate that *PpMS1A* and *PpMS1B* have functions also in the diploid sporophyte generation. Their expression domains suggests that *PpMS1* activity is dispensable for early embryo development but is likely to have functions in transfer cells of the sporophyte foot and in the developing sporangium. In the sporangium, the *PpMS1* genes are primarily expressed in the sporogenous cell layer soon after its establishment, supporting a possible function of *PpMS1* activity for the specification of these cells. In angiosperms, MS1 and MMD1 are part of a complex gene regulatory network facilitating tapetum-mediated pollen development (Ferguson *et al*., 2017; Lei & Liu, 2020). Among other factors, this network also includes bHLH clade II and III(a+c) genes. The PpMS1 expression pattern in moss sporophytes, combined with that of PpbHLH clade II and III(a+c) genes (Lopez-Obando *et al*., 2022), support that moss sporogenesis could be regulated by a homologous network inherited from the common ancestor of land plants.

Our finding that *PpMS1A* and *PpMS1B* provide functions in sporophytes as well as during male and female gametogenesis, while function of their angiosperm homologs is restricted to processes finally leading to completion of male gametogenesis only, raises questions about what the original PHD clade IIa function in the ancestors of all extant land plants may have been. We speculate that the functions in antheridia, archegonia and sporophytes of mosses today all may originate from one and the same function in a hypothetical ancestral plant, with morphologically similar gametes (isogamy) and a haplontic life cycle where the zygote underwent meiosis without intervening mitotic divisions. Thus, we hypothesize that PHD clade IIa activity served to specify the two gametes, where after it was carried over via fertilization to the zygote, in which it secured expression of genes needed both for nutrient uptake from the gametophyte generation and for meiosis. Later during evolution, when mitotic divisions of the zygote morphologically separated the cells destined for meiosis from the cells carrying over nutrients from the gametophyte, this was accompanied by a similar spatiotemporal separation of clade IIa activity.

To challenge this speculative hypothesis, it would be highly interesting to investigate the expression pattern and function of possible PHD clade IIa homologs in the algal sisters of land plants (Ito *et al*., 2007). Using a recent transcriptome study (Sanchez-Vera *et al*., 2022), we in fact identified 135 additional moss genes with significantly higher expression in egg cells, antheridia and the diploid sporophyte generation at the green stage when sporogenesis occur, compared to vegetative haploid tissues (Table S4). They thus have the potential to play specific roles in both gametogenesis and sporogenesis. Possibly, the hypothetical evolutionary history outlined for clade IIa genes could be shared also with these genes. They include the two *PpBNB* genes encoding class VIIIa bHLH transcription factors. While the Arabidopsis *BNB* genes are required only for specification of male generative cells, the *Marchantia polymorpha* homologs are essential for the initiation of both male and female reproductive organs, and are active in egg and sperm progenitors (Yamaoka *et al*., 2018; Hisanaga *et al*., 2019). In *P. patens, BNB’s* are important for both male and female germ cell specification, making it difficult to assess their potential role in the green sporophyte, where it is also expressed (Sanchez-Vera *et al*., 2022). A moss gene, *PpMKN1*, encoding a class 2 KNOTTED1-LIKE HOMEOBOX (KNOX2) transcription factor preventing haploid-specific development in the sporophyte phase (Sakakibara *et al*., 2013) is also significantly up-regulated in the egg and antheridia, although at a much-reduced level compared to the green sporophyte. Additional transcription factor genes, such as homologs to Arabidopsis *WRI2*, *LEC2, EFM, NAC56*, are also up-regulated in all three reproductive tissue types, but their functions in moss are unknown and not easy to extrapolate from their functions in Arabidopsis. A homolog to CCR4-NOT complex component NOT1, which in Arabidopsis regulates RNA-directed methylation and transcriptional silencing (Zhou *et al*., 2020), is also elevated in the reproductive organs and the green sporophytes. AtNOT1 is necessary for e.g. proper male germ cell development, pollen germination and embryogenesis (Motomura *et al*., 2020; Pereira *et al*., 2020). A moss homolog of MBD9, a SWR1-C interacting protein required for H2A.Z deposition at a subset of actively transcribing genes in Arabidopsis (Potok *et al*., 2019; Luo *et al*., 2020), also show elevated expression in the three selected tissue types.

Assuming that the need for PHD clade IIa functions in moss to complete male and female gametogenesis as well as sporogenesis represent a heritage from ancestral land plants, one can ask how this has evolved into the clade IIa-regulation in anthers evident in angiosperms. It appears very likely that this can be attributed to the loss of gametangia as part of a dramatic reduction of the gametophyte generation in angiosperms (Hisanaga *et al*., 2019). Thus, the need for clade IIa-functions during gametogenesis may either have been lost, or at least become difficult to separate from sporophytic functions facilitating meiosis, as the two processes have become so intimately coupled in time and space in angiosperms.

The possible homology between clade IIa functions in angiosperm anthers and moss sporangia is complicated by the fact that Arabidopsis *AtMS1* and *AtMMD1* exert their functions in distinct anther cell types. Thus, *AtMMD1* controls gene expression and chromosome condensation in microsporocytes (Yang *et al*., 2003; Reddy *et al*., 2003) while *AtMS1* controls gene expression in tapetal cells (Wilson *et al*., 2001; Ito & Shinozaki, 2002; Alves-Ferreira *et al*., 2007; Yang *et al*., 2007; Reimegård *et al*., 2017; Lu *et al*., 2020). Our phylogenetic analysis reveals that bryophyte clade IIa homologs cluster with angiosperm MS1 proteins rather than with MMD1 proteins. While this could indicate that the duplication event giving rise to *MS1* and *MMD1* took place already in the common ancestors of all extant land plants but that *MMD1* homologs have been lost in non-seed plant lineages through the course of evolution, we find it equally likely that the duplication event took place in the angiosperm lineage after its divergence from bryophytes where after MMD1-clade genes diversified by neo- or sub-functionalization.

Even if the current study does not address the molecular basis of *PpMS1* activity, the conservation of key protein domains like the N-terminal nuclear localization signal, the central MMD domain, and the C-terminal PHD domain supports that also bryophyte PHD clade IIa proteins functions by controlling cell identity through the regulation of chromatin structure and gene expression (Andreuzza *et al*., 2015; Reimergård *et al*., 2017; Wang *et al*., 2020). Future studies will have to reveal if this is indeed the case, and to what extent related genes and gene clusters are targeted by clade IIa-regulation in the angiosperm anther and in the reproductive organs and the sporophyte of moss.

## Acknowledgments

We thank Ulf Lagercrantz for help with bioinformatics analyses on which Table S4 is based on. This work was supported by grants from the Swedish Research Council to ES and MT (621-2014-4941; 2018-04068) and the Nilsson-Ehle Endowments to KL and MLO.

## Author contributions

KL, MLO, VSV and MT conducted the experimental work and analyzed the data. MLO, KL, ES and MT designed experiments and interpreted data. MT, ES and KL wrote the manuscript.

## Data availability

The data that support the findings of this study are available from the corresponding author upon request.

## Supporting Information

Additional Supporting Information may be found online in the Supporting Information tab for this article:

**Figure S1** Overviews of how reporter lines were generated.

**Figure S2** Overviews of how *PpMS1A* and *PpMS1B* loss-of-function mutants were generated.

**Figure S3** Data from main figure 1c-d presented in a way making comparisons of transcript abundance between the two genes possible.

**Figure S4** Amino acid sequence alignment used to infer phylogenetic tree in Fig. 1.

**Figure S5** Non-filtered full length alignment of all proteins in Fig. 1 belonging to the MS1- and MMD1-clades.

**Figure S6** Confocal microscopy images showing *PpMS1B::GFPGUS-1* GFP reporter signals in sporophytes.

**Table S1** Primers used in this study.

**Table S2** Characteristics of knock-out and knock-in lines obtained by CRISPR-CAS9 gene editing.

**Table S3** Characteristics of crRNAs in gRNA-expressing plasmids.

**Table S4** *P. patens* genes for which publically available RNA-seq data indicates higher expression in green sporophytes, eggs and antheridia, respectively, than in vegetative tissue samles. – see separate excel file

## Notes

### Competing Interest Statement

The authors have declared no competing interest.

