## Supplemental Information for "*MS1/MMD1* homologs in the moss *P. patens* are required for male and female gametogenesis and likely for sporogenesis"

*New Phytologist* **Supporting Information**

Article acceptance date: Click here to enter a date.

The following Supporting Information is available for this article:

**Figure S1** Overviews of how reporter lines were generated.

**Figure S2** Overviews of how *PpMS1A* and *PpMS1B* loss-of-function mutants were generated.

**Figure S3** Data from main figure 1c-d presented in a way making comparisons of transcript abundance between the two genes possible.

**Figure S4** Amino acid sequence alignment used to infer phylogenetic tree in Fig. 1.

**Figure S5** Non-filtered full length alignment of all proteins in Fig. 1 belonging to the MS1- and MMD1-clades.

**Figure S6** Confocal microscopy images showing *PpMS1B::GFPGUS-1* GFP reporter signals in sporophytes.

**Table S1** Primers used in this study.

**Table S2** Characteristics of knock-out and knock-in lines obtained by CRISPR-CAS9 gene editing.

**Table S3** Characteristics of crRNAs in gRNA-expressing plasmids.

**Table S4** *P. patens* genes for which publically available RNA-seq data indicates higher expression in green sporophytes, eggs and antheridia, respectively, than in vegetative tissue samles – see separate excel file

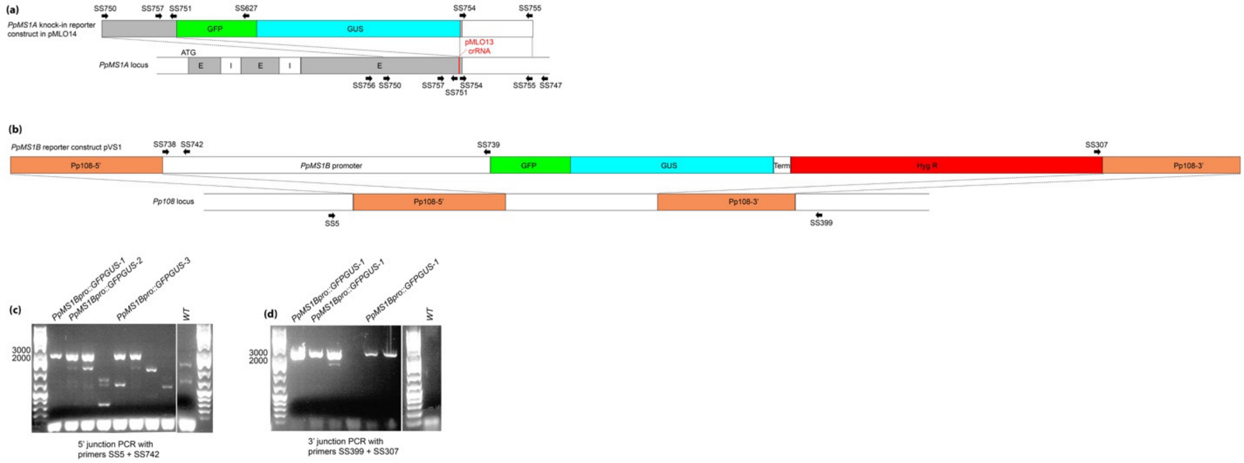


Fig. S1. Overviews of how reporter lines were generated. (a) Schematic view of the *PpMS1A* translational reporter construct pMLO14 (top) and the *PpMS1A* locus into which it was inserted (bottom). Annealing sites of primers used to build the construct and to confirm resulting lines by PCR amplification and sequencing are shown as arrows. The target site of the pMLO13 crRNA, used to increase the frequency of correct targeting, is indicated in red. (b) Schematic view of the *PpMS1B* transcriptional reporter construct pVS1 (top) and the neutral *Pp108* locus to which it was targeted (bottom). Annealing sites of primers used to build the construct, and to confirm resulting lines by PCR amplification in (c) and (d) are shown as arrows. (c,d) PCR verification of 5’ (c) and 3’ (d) junctions to confirm correct integration in lines transformed with the *PpMS1B* reporter construct pVS1. Annealing sites of primers used are indicated in (b), and the sequences of primers used are shown in Table S1.

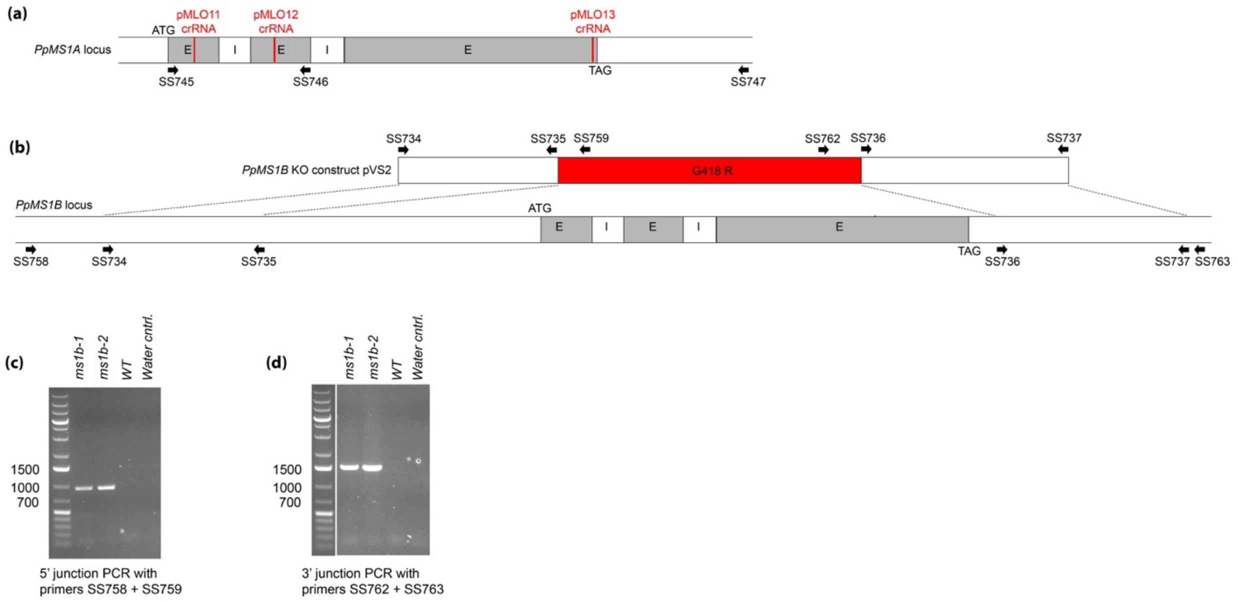


Fig. S2. Overviews of how *PpMS1A* and *PpMS1B* loss-of-function mutants were generated. (a) Schematic view of the *PpMS1A* locus indicating the target sites of cRNAs used to generate CRISPR-CAS9 loss-of-function mutants (red marks) and the annealing sites of primers used to evaluate the gene editing outcome by PCR amplification and sequencing (arrows). (b) Schematic view of the *PpMS1B* knockout construct pVS2 (top) and the *PpMS1B* locus (bottom) to which it was targeted. Annealing sites of primers used to build the construct, and to confirm resulting lines by PCR amplification in (c) and (d) are shown as arrows. (c,d) PCR verification of 5' (c) and 3' (d) junctions to confirm correct integration in lines transformed with the *PpMS1B* knockout construct pVS2. Annealing sites of primers used are indicated in (b), and the sequences of primers used are shown in Table S1.

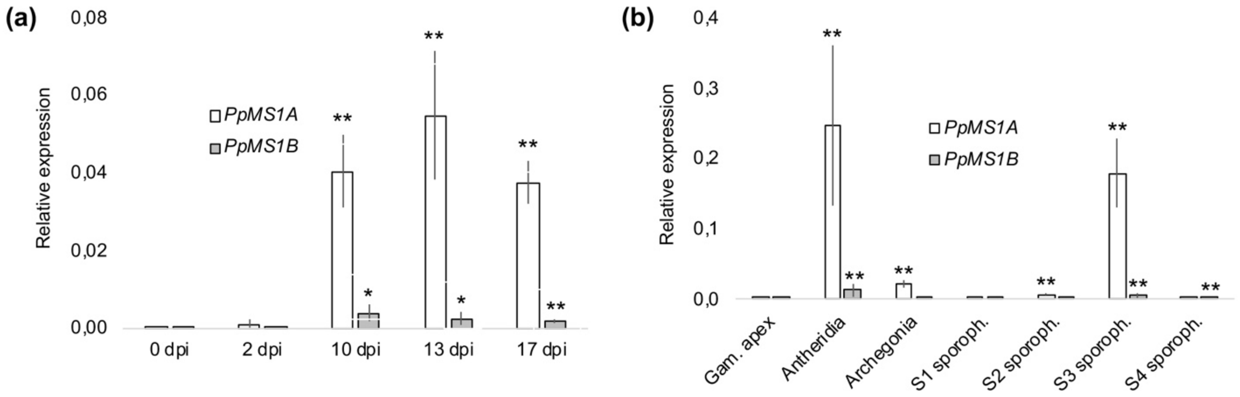


Fig. S3. Data from main figure 1c-d presented in a way making comparisons of transcript abundance between the two genes possible. (a) Relative transcript abundance of *PpMS1A* and *PpMS1B* in WT gametophore apices at different days post induction (dpi) of reproductive development. Typical occurrence of reproductive organs at selected time points: 0 and 2 dpi, no reproductive organs; 10 dpi, young antheridia; 13 dpi, mid-stage antheridia and young archegonia; 17 dpi, mature antheridia and mid-stage archegonia. (b) Relative transcript abundance of *PpMS1A* and *PpMS1B* in isolated WT gametophore shoot apices (without reproductive organs), antheridia bundles, archegonia bundles, and sporophytes of different developmental stages. Sporophyte samples contain organs roughly correlating to the following stages described in Lopez-Obando et al. (2022): S1, st.1-3; S2, st.4-8; S3, st.9-11; S4, st.12-14. In both (c) and (d), the sample with the highest transcript abundance regardless of whether *PpMS1A* or *PpMS1B* is considered was set to 1, each data point represents an average of three independent biological replicates, error bars indicate standard deviation and asterisks indicate a statistically significant difference from gametophore apex sample prior to reproductive organ formation (Student´s t-test: *, $P < 0.05$; **, $P < 0.02$).

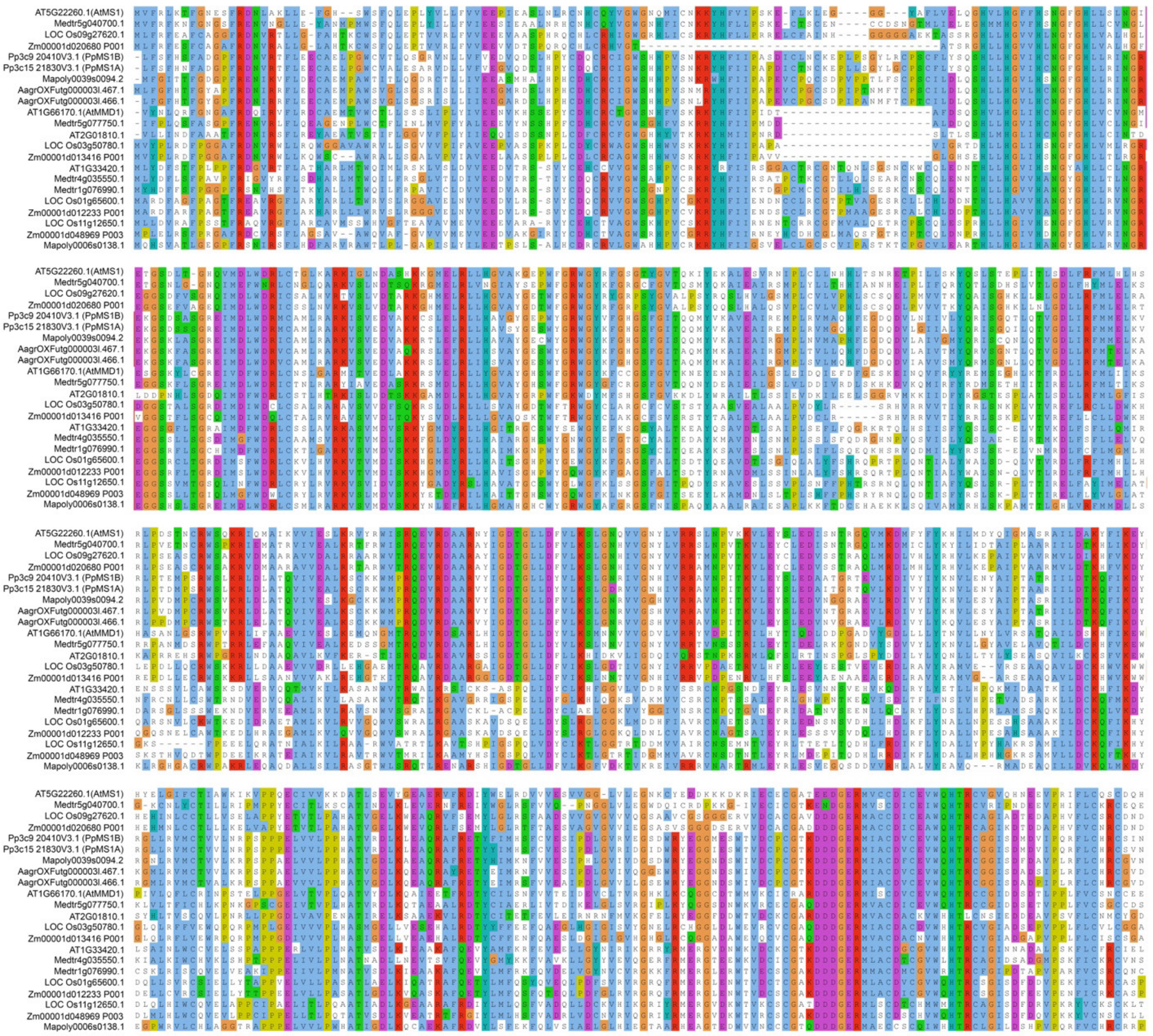

Fig. S4. Amino acid sequence alignment used to infer phylogenetic tree in Fig. 1. Similar amino acids are marked by the same color. Note that the alignment was filtered using Transitive Consistency Scores (Chang et al., 2014) to facilitate accurate phylogenetic reconstruction. At: *Arabidopsis thaliana* (dicot), Medtr: *Medicago truncatula* (dicot), LOC Os: *Oriza sativa* (monocot), Zm: *Zea mays* (monocot), Pp: *Physcomitrium patens* (moss), AagrOXF: *Anthoceros agrestis* (Oxford; hornwort), Mapoly: *Marchantia polymorpha* (liverwort).

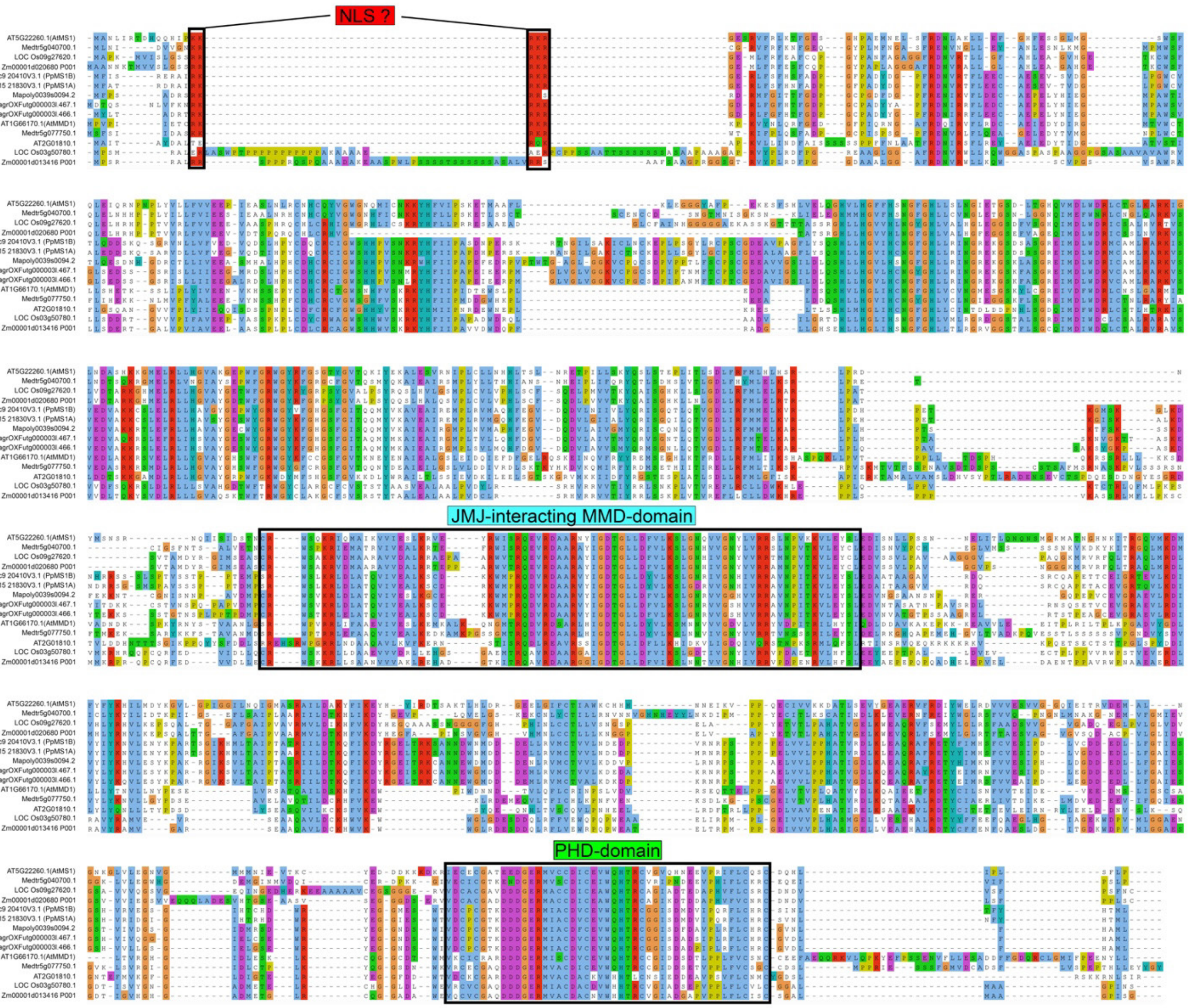


Fig. S5. Non-filtered full length alignment of all proteins in Fig. 1 belonging to the MS1- and MMD1-clades. Similar amino acids are marked by the same color. Conserved domains that have been assigned possible functions in flowering plants are boxed: Red, possible nuclear localization signal (NLS); Blue, JMJ-interacting domain; Green, PHD domain. At: *Arabidopsis thaliana* (dicot), Medtr: *Medicago truncatula* (dicot), LOC Os: *Oriza sativa* (monocot), Zm: *Zea mays* (monocot), Pp: *Physcomitrium patens* (moss), AagrOXF: *Anthoceros agrestis* (Oxford; hornwort), Mapoly: *Marchantia polymorpha* (liverwort).

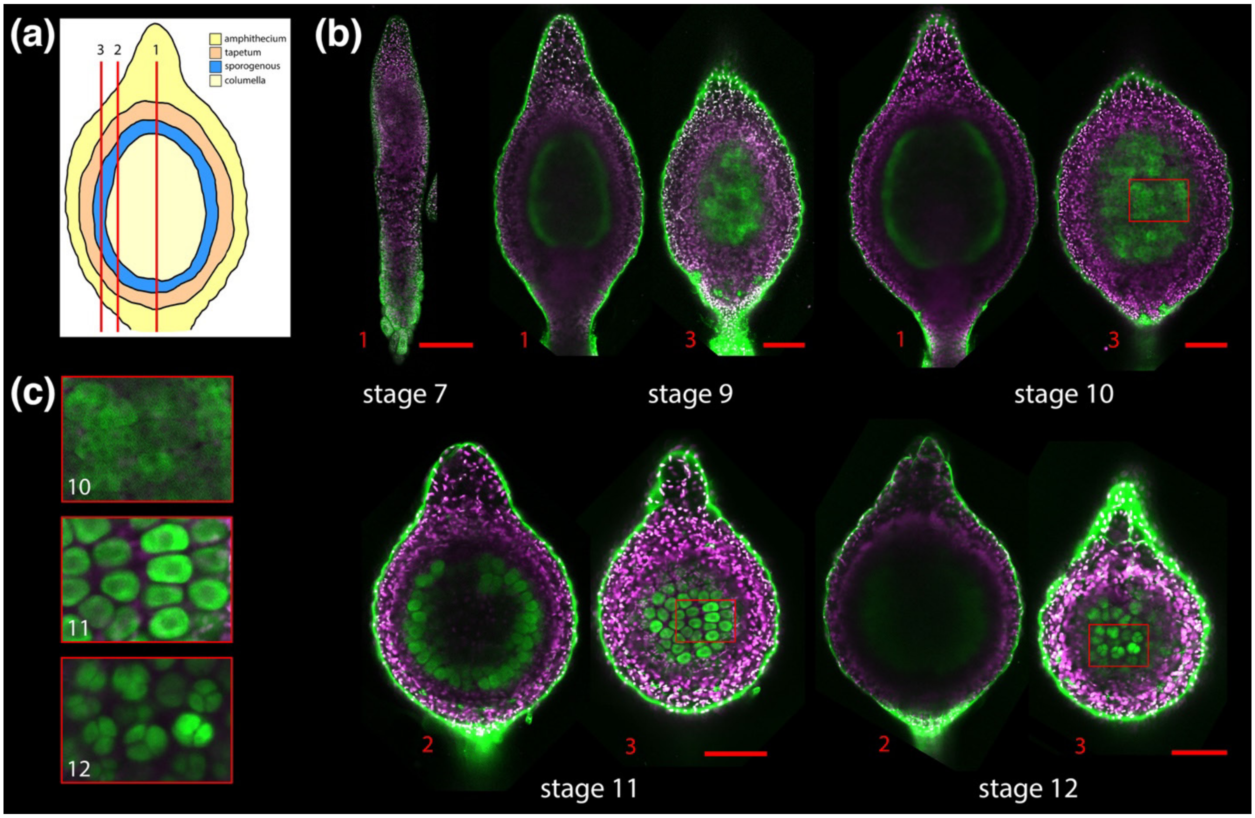


Fig. S6. Confocal microscopy images showing *PpMS1B::GFPGUS-1* GFP reporter signals in sporophytes. (a) Principle sporangium sketch representative for stage 9-12 showing main cell layers (for details, see Lopez-Obando et al., 2022) and red lines indicating the approximate focal planes of images in (b). (b) Typical *PpMS1B::GFPGUS-1* GFP reporter signals in a stage 7 sporophyte and in stage 9, 10, 11 and 12 sporangia. Each image shows a merge of confocal channels detecting green fluorescent protein (green) and chloroplast autofluorescence (magenta). Numbers in red indicates the approximate focal plane as outlined in (a). Note signals in transfer cell of foot at stage 7 and in sporogenous cells in stage 9-12. The green rim lining the outer contour of organs represents an optical artifact rather than an actual GFP signal. Size bars, 100 µm. (c) Boxed areas with sporogenous cells in stage 10, 11 and 12 organs in (b) shown at 3 times magnification.

**Table S1.** Primers used in this study

| Name | Sequence (5' to 3') |
|---|---|
| SS5 | GAAACCTCCCAAGCTCTGACGA |
| SS307 | CGTCCGAGGGCAAAGAAATAGAGTA |
| SS399 | TGCACCTCATCGACCCTTCAGT |
| SS584 | GTCGATGCATTGGGTGGAGTCA |
| SS585 | CTCTTTGAGCGCTCGGGATTGT |
| SS586 | GCAATGCTGAGAGCCAGGAAAGTC |
| SS587 | GTCCGAATTTGTATCCCCACCTTC |
| SS611 | GGGGACAACTTTTCTATACAAAGTTGAATTCCCATGGAGTCAAAG |
| SS612 | GGGGACAACTTTATTATACAAAGTTGTAATTCGAGCTCGGTACCCAC |
| SS627 | TCAGGGCGGACTGGGTGCT |
| SS734 | GGGGACAAGTTTGTACAAAAAAGCAGGCTCCTAGGGGTCACTGCACAAAATGGGT |
| SS735 | GGGGACAACTTTGTATAGAAAAGTTGGGTGTCCTGGTCAGTGGAACCG |
| SS736 | GGGGACAACTTTGTATAATAAAGTTGCTTTTCGTGGCCCAGAGAAG |
| SS737 | GGGGACCACTTTGTACAAGAAAGCTGGGTAGTTAACATAGTTGACCCTGGAGCCAC |
| SS738 | TAAGGATCCGAGACTAACCTTGAAAATAATAC |
| SS739 | TGACCATGGCTGCCAATAGGTGAGCATTTA |
| SS742 | GCTCTAAATAAGACAGCATATGATGGC |
| SS743 | CCATGGAGGTTAACGCTGCATCGA |
| SS744 | AAACTCGATGCAGCGTTAACCTCC |
| SS745 | GAGATCGAGCAATCAGAAAGC |
| SS746 | AAAGGTCCATGATCTCCCTTC |
| SS747 | CCCGTTGTATGAGGGTTCCTTC |
| SS748 | ACATAAGCCCTCGTAACAAAGCCCGAAAGGAAGCTGAG |
| SS749 | CAACTTCACAGGCAACCGTTGTGGTCTCCCTATAGT |
| SS750 | GACCACAACGGTTGCCTGTGAAGTTGGAAGGACC |
| SS751 | CCTTGCTCACCATCAGGAAGCGTTGAGGAATTACATCC |
| SS752 | TCAACGCTTCCTGATGGTGAGCAAGGGCGAG |
| SS753 | GGTTAACGCTGCATCATTGTTTGCCTCCCTGCT |
| SS754 | AGGCAAACAATGATGCAGCGTTAACCTCAATTTCTATCAC |
| SS755 | TCGGGCTTTGTTACGAGGGCTTATGTTTTCTGCGA |
| SS756 | ACTATGTCCTGAAATCCTTGGGT |
| SS757 | ATCCATACTCGTCATGACTGGC |
| SS758 | TAATACCCGTTTGCAAGTGAGTCA |
| SS759 | CTTTACGGCGAGTTCTGTTAGGTC |
| SS762 | GCCTTCTATCGCCTTCTTGACG |
| SS763 | CATCCTGAAACTGTGTCCACTGGT |

**Table S2.** Characteristics of knock-out and knock-in lines obtained by CRISPR-CAS9 gene editing.

| Line name | Gene: gRNA plasmids | Mutation pattern[a] | Predicted consequence |
|---|---|---|---|
| *ms1a-1* | *PpMS1A*: pMLO11 + pMLO13 | WT: AGTCACAGTCGATGGCTTACC 2216b CAACGCTTCCTGTGCCATCGA<br>M: AGTCACAGTCGATGGCTT----------------- CCTGTGCCATCGA | Knock-out, protein truncated at aa 74 (of 691) |
| *ms1a-2* | *PpMS1A*: pMLO11 + pMLO13 | WT: AGTCACAGTCGATGGCTTACC 2216b CAACGCTTCCTGTGCCATCGA<br>M: AGTCACAGTCGATGG---------------------------CCATCGA | Knock-out, protein truncated at aa 72 (of 691) |
| *ms1a-3* | *PpMS1A*: pMLO11 + pMLO12 | WT: AGTCACAGTCGATGGCTTACC 429b TTACTATCAGGCTACCTCGG<br>M: AGTCACAGTCGACTTTATA------------TATCAGGCTACCTCGG | Knock-out, protein truncated at aa 125 (of 691) |
| *ms1ams1b-1* | *msa1b-1* deletion | WT:<br>M: | Knock-out |
|  | *PpMS1L2*: (pMLO11 + pMLO12) | WT: AGTCACAGTCGATGGCTTACC 429b TTACTATCAGGCTACCTCGG<br>M: AGTCACAGTCGA------------------CTATCAGGCTACCTCGG | Knock-out, protein truncated at aa 123 (of 691) |
| *ms1ams1b-2* | *msa1b-1* deletion | WT:<br>M: | Knock-out |
|  | *PpMS1A*: (pMLO11 + pMLO12) | WT: AGTCACAGTCGATGGCTTACC 429b TTACTATCAGGCTACCTCGG<br>M: AGTCACAGTCGATGGCT---------------ATCAGGCTACCTCGG | Knock-out, protein truncated at aa 124 (of 691) |
| *PpMS1Apro::PpMS1A-GFPGUS-1* | *PpMS1A*: (pMLO13 + pMLO14) | WT: TGTAATTCCTCAACGCTTCCTGTGCCATCGA<br>M: TGTAATTCCTCAACGCTTCCTG---------GFPGUS | Knock-in, 676 aa of PpMS1A fused in frame to GFPGUS |
| *PpMS1Apro::PpMS1A-GFPGUS-2* | *PpMS1A*: (pMLO13 + pMLO14) | WT: TGTAATTCCTCAACGCTTCCTGTGCCATCGA<br>M: TGTAATTCCTCAACGCTTCCTG---------GFPGUS | Knock-in, 676 aa of PpMS1A fused in frame to GFPGUS |
| *PpMS1Apro::PpMS1A-GFPGUS-3* | *PpMS1A*: (pMLO13 + pMLO14) | WT: TGTAATTCCTCAACGCTTCCTGTGCCATCGA<br>M: TGTAATTCCTCAACGCTTCCTG---------GFPGUS | Knock-in, 676 aa of PpMS1A fused in frame to GFPGUS |

a: Protospacer adjacent motif (PAM) sequence is underlined, 20 bases of the crRNAs are colored in red, deletions are indicated by dashes, insertions are highlighted in yellow.

**Table S3.** Characteristics of crRNAs in gRNA-expressing plasmids.

| Plasmid name | Use | crRNA target | crRNA sequence and SS (%)[a] | Predicted off-target sequences[b] |
|---|---|---|---|---|
| pMLO11 | Knock-out | *PpMS1A* (res. 157 - 179 of CDS) | GTCACAGTCGATGGCTTACC*CGG* (100) | GTCACAGTCGAAGGATGCCC<br>GTCACAGTCGACGTCTGACA<br>GTTTCCGTCGATGGCCTACC |
| pMLO12 | Knock-out | *PpMS1A* (res. 421 – 443 of CDS) | CCGAGGTAGCCTGATAGTAACGG (100) | GCGAGGCAGCCTGATATTGA |
| pMLO13 | Knock-out and Knock-in | *PpMS1A* (res. 2034 - 2053 of CDS) | TGAGGTTAACGCTGCATCGA*TGG* (98) | TGATGTTGATGCAGCATCGA<br>TAAGGATAAAGCTGCGTCGA<br>TGGGGTGCTCGCTGCATCGA<br>AGAGGTTAACTCTACAGCGA<br>AGAGGTTAACACTGCAGGGA<br>GGAGGTTCACGCTGCGGCGA<br>TGACGTGGACGCTGCCTCGA<br>TGAGGTTAAACCTGATTCGA |

a: Protospacer adjacent motif (PAM) sequence preceded by crRNA sequence is shown in italics and underlined. SS; Specificity score from CRISPOR (Haeussler et al., 2018)

b: Mismatch of off-target sequence with respect to the crRNA sequence is shown in red
